# Centromere-Proximal Suppression of Meiotic Crossovers in *Drosophila* is Robust to Changes in Centromere Number and Repetitive DNA Content

**DOI:** 10.1101/2023.10.17.562696

**Authors:** Nila M. Pazhayam, Leah K. Frazier, Jeff Sekelsky

## Abstract

Accurate segregation of homologous chromosomes during meiosis depends on both the presence and regulated placement of crossovers (COs). The centromere effect (CE), or CO exclusion in pericentromeric regions of the chromosome, is a meiotic CO patterning phenomenon that helps prevent nondisjunction (NDJ), thereby protecting against chromosomal disorders and other meiotic defects. Despite being identified nearly a century ago, the mechanisms behind this fundamental cellular process remain unknown, with most studies of the *Drosophila* CE focusing on local influences of the centromere and pericentric heterochromatin. In this study, we sought to investigate whether dosage changes in centromere number and repetitive DNA content affect the strength of the CE, using phenotypic recombination mapping. Additionally, we also studied the effects of repetitive DNA function on CE strength using satellite-DNA binding protein mutants shown to have defective centromere clustering. Despite what previous studies suggest, our results show that the *Drosophila* CE is robust to dosage changes in centromere number and repetitive DNA content, and potentially also to repetitive DNA function. Our study suggests that the CE is unlikely to be spatially controlled, providing novel insight into the mechanisms behind the *Drosophila* centromere effect.

## Introduction

Meiosis is a specialized form of cell division in which a diploid chromosome set is reduced to a haploid set through the segregation of homologous chromosomes during anaphase of meiosis I (MI). Accurate disjunction, or segregation, is largely facilitated by crossing-over between homologs, an integral part of MI that requires the formation of programmed double-strand breaks (DSBs) along the chromosome and involves reciprocal exchange of chromosome arms. DSBs are repaired through homologous recombination to generate either crossover (CO) or noncrossover (NCO) repair products. The decision of whether a DSB is repaired as a CO or NCO is a highly regulated process, as the frequency and positioning of meiotic COs is critical for proper segregation of homologs.

A far greater number of DSBs are formed during meiosis than are repaired as COs and the mechanisms that regulate CO placement along the chromosome are collectively referred to as CO patterning phenomena. Three important patterning events are: 1) assurance, which dictates that every pair of homologs receives at least one CO (OWEN 1950), 2) interference, which ensures that COs do not form right next to one another (STURTEVANT 1913) and 3) the centromere effect (CE), which dictates that COs are excluded from centromere-proximal regions (BEADLE 1932). These have been reviewed extensively in (PAZHAYAM *et al*. 2021). These phenomena are critical for accurate disjunction of homologous chromosomes in humans, safeguarding against miscarriages and chromosomal disorders.

The centromere effect in particular has shown to be vital for preventing non-disjunction (NDJ), the risk of which increases with increasing maternal age. Studies in both humans and *Drosophila melanogaster* have established a direct link between pericentromeric crossing over and NDJ [(LAMB *et al*. 1996) (KOEHLER *et al*. 1996)]. Furthermore, human studies have established that mis-segregation events on chromosome *21*, the leading cause of Down syndrome, correlate with centromere-proximal COs and increase with maternal age (OLIVER *et al*. 2012). However, very little is known about the mechanisms behind the CE, despite it being a critical safeguard against chromosome mis-segregation and age-associated meiotic defects.

The CE was first reported in 1932 when Beadle observed a decrease in CO frequencies in *D*. *melanogaster* translocation stocks that had a portion of the third chromosome moved closer to the centromere of the fourth than it had originally been to the centromere of the third (BEADLE 1932). He attributed this reduction in recombination rate to the chromosomal interval’s increased proximity to the chromosome *4* centromere. Although first described over nine decades ago, the mechanisms behind the CE have remained elusive to this day.

In most organisms, centromeres are surrounded by pericentromeric heterochromatin, consisting of repetitive DNA that includes large arrays of satellite DNA along with other repetitive elements. In *D. melanogaster*, nearly one-third of the genome is heterochromatic pericentromeric satellite DNA (HOSKINS *et al*. 2002). A prominent question regarding the mechanism of the CE has centered on the ways in which this heterochromatic, repetitive sequence and the centromere itself contribute to centromere-proximal CO suppression. Of the handful of studies that have addressed pericentromeric crossing over in the past century, most have attempted to establish one as more important than the other in *D. melanogaster*, with contrasting results.

Heterochromatin has been considered everything from an active participant in CO reduction in adjacent intervals [(SLATIS 1955), (JOHN 1985), (WESTPHAL AND REUTER 2002), (MEHROTRA AND MCKIM 2006)] to nothing more than a passive spacer [(MATHER 1939), (YAMAMOTO AND MIKLOS 1978a)] between euchromatin and the centromere.

The influence of repetitive DNA on *D*. *melanogaster* CO frequencies has been studied in the past, however always in *cis.* A 1955 study measured COs in homozygous *bw^D^* mutants, flies with an ∼2Mb insertion of {AAGAG}_n_ satellite sequence into the distal *brown* locus on chromosome *2R* and observed a marked reduction in CO frequencies surrounding the insertion (SLATIS 1955). The exclusion of DSBs in heterochromatic regions of flies, which are primarily repetitive, has also been shown (MEHROTRA AND MCKIM 2006), suggesting that CO suppression in pericentromeric regions may be explained by decreases in DSB formation in the region. Decreased dosage of chromosome *4,* which consists almost entirely of repetitive sequence, has also been shown to have *cis*-effects, reducing the expression of various chromosome *4* genes (HAYNES *et al*. 2007).

Furthermore, repetitive DNA has been shown to have *trans*-effects on phenomena such as gene expression and heterochromatic integrity in *D*. *melanogaster*. In this study, we use the term *trans* to refer to effects on other chromosomes and not on the other homolog. Flies with extra copies of chromosome *Y*, which consists almost entirely of highly-repetitive DNA, have been shown to de-repress position effect variegation, the phenomenon where proximity to heterochromatin causes variegated gene expression [(MULLER 1930), (DIMITRI AND PISANO 1989)]. This is thought to be due to the *Y* chromosome increasing competition for a limited pool of heterochromatic factors within a cell, leading to a decrease in heterochromatinization elsewhere in the genome, and the consequent de-repression of genes close to heterochromatic boundaries. Several recent studies support this proposal, with lowered di- and trimethylation of H3K9, a canonical mark of heterochromatin, observed at the pericentromeres of all chromosomes in *XXY* and *XYY* flies, compared to *XX* and *XY* flies (BROWN *et al*. 2020), and increasing *Y* chromosome lengths negatively correlating with gene silencing in *trans* (DELANOUE *et al*. 2023). Despite the implications of these studies, a key piece of the puzzle that remains unanswered is whether repetitive DNA and heterochromatin exert *trans*-effects on meiotic CO frequencies, particularly near the centromere.

*Cis*-effects of the centromere on CO frequencies have also been established in *D*. *melanogaster*, the first instance of which was Beadle concluding that CO frequencies in euchromatic regions decrease when brought closer to a centromere (BEADLE 1932). In his 1939 study, Mather further concluded that CO frequencies in euchromatin depended more on proximity to the centromere than on proximity to heterochromatin (MATHER 1939). Similarly, Yamamoto and Miklos moved euchromatic regions closer to the centromere through deleting pericentromeric heterochromatin, showing that CO frequencies in these regions negatively correlate with proximity to the centromere (YAMAMOTO AND MIKLOS 1978b). In the 1930s, Helen Redfield measured COs in triploid *D. melanogaster* females, and observed substantial increases in centromere-proximal CO frequencies, compared to diploids [(REDFIELD 1930), (REDFIELD 1932)]. Although this change in CE strength may be a result of ploidy changes, it is also possible that it is a consequence of the total number of centromeres in triploids increasing 1.5-fold, raising key questions about whether dosage changes in just centromere number are capable of exerting *trans*-effects on centromere-proximal CO frequencies.

To fill this gap in knowledge regarding the mechanisms of a fundamental cellular process, we measured centromere-proximal CO-frequencies and CE strength in mutants with dosage changes in centromere number and repetitive DNA content, respectively. We also measured centromere-proximal CO frequencies in mutants of satellite-DNA binding proteins to ask if repetitive DNA function plays a role in the establishment of the CE. Surprisingly, our results show no change in the strength of the CE in mutants with a decreased total number of centromeres, mutants with increased and decreased total repetitive DNA content, or with loss-of-function mutants of satellite DNA-binding proteins. Overall, our study suggests that the CE is robust to dosage changes in centromere number and amount of repetitive DNA, and presumably also repetitive DNA function.

## Results

### Centromere dosage does not exert *trans*-effects on the CE

While the centromere’s local, cis-acting effect on CO frequencies in centromere-proximal chromosomal regions has been established in *D. melanogaster* (YAMAMOTO AND MIKLOS 1978a), whether centromeres also exert *trans*-effects on CO frequencies remains unknown. Although Redfield showed that triploid *D. melanogaster* females have an increased number of centromere-proximal COs on chromosomes *2* and *3* compared to diploids [(REDFIELD 1930) (REDFIELD 1932)], it is unclear whether this effect is due to a change in total centromere number from 16 to (one per sister chromatid for eight chromosomes) to 24 in triploids (one per sister chromatid for twelve chromosomes), or a consequence of ploidy changes. We hypothesized that since Redfield’s experiments showed a weakened CE with an increase in centromere numbers, a reduction in centromeres would lead to a strengthened CE. To investigate the effects of reducing total centromere number without the complicating factor of ploidy change, we made use of *D. melanogaster* stocks with compound *X* and *4th* chromosomes. Flies of this genotype have attached homologs of chromosome *X* as well as attached homologs of chromosome *4*, with each attached chromosome only having two centromeres, thereby reducing total centromere number from 16 to 12 (Fig. 1A).

**Figure 1.**
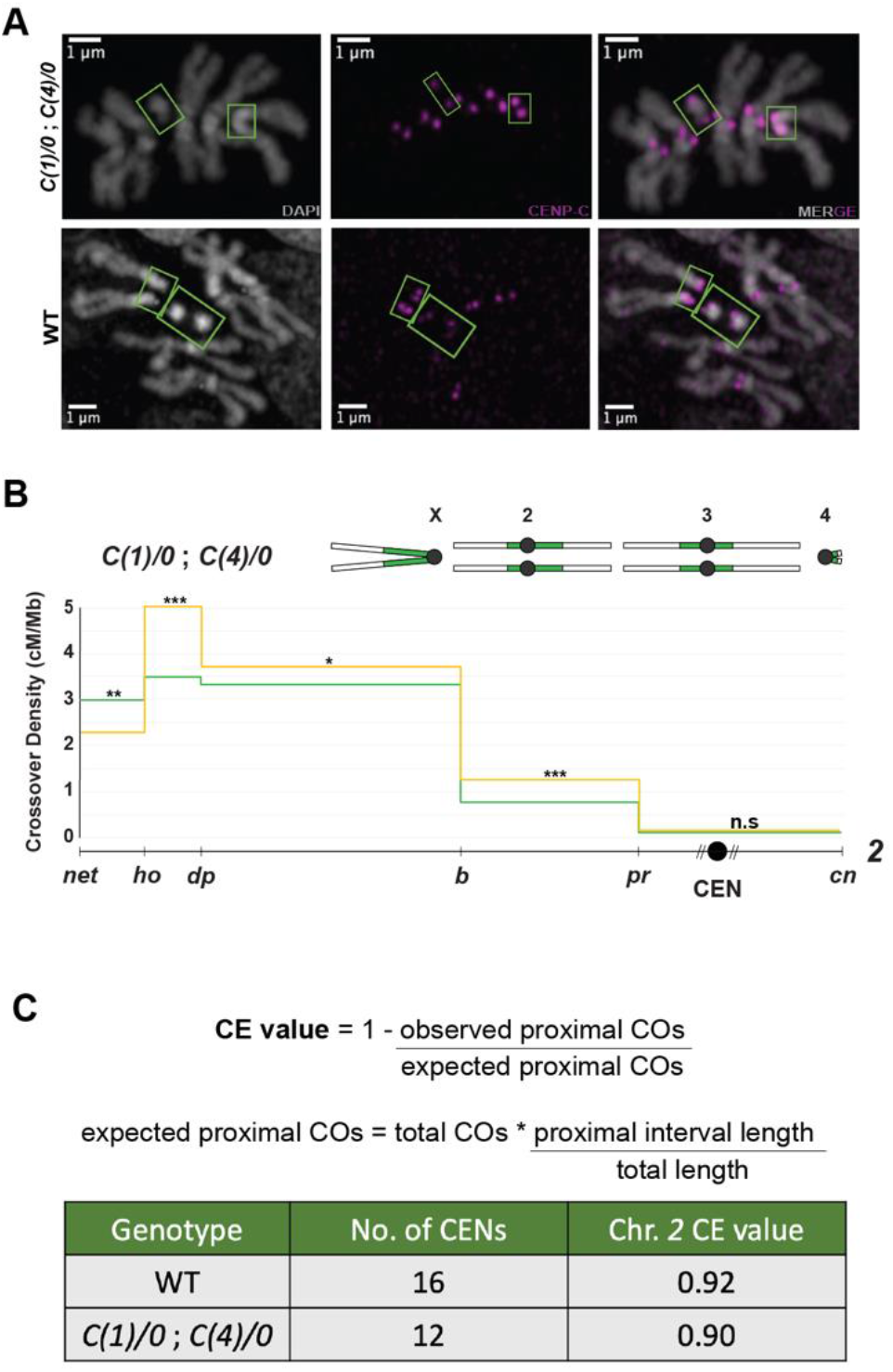
Centromere dosage does not exert *trans*-effects on the CE. **(A)** Mitotic chromosome spreads (grey) with CENP-C foci (magenta) from *C(1)/0*; *C(4)/0* larvae (top panel) and wild type larvae (bottom panel). Green boxes highlight chromosomes *X* (larger, V-shaped structure) and *4* (smaller, dot-like structure) in both genotypes. **(B)** CO distribution along chromosome *2* in *C(1)/0*; *C(4)/0* flies (yellow line, *n*=5311) and wild type flies (green line, *n*= 4104), with a schematic of the *C(1)/0*; *C(4)/0* karyotype shown above the graph. CO density in cM/Mb is indicated on the Y-axis, relative physical distances between markers used to score COs are indicated on the X-axis. The black circle represents the centromere, with dashed lines around it representing pericentromeric repetitive sequence that remains unassembled. Statistical significance in each interval was calculated using two-tailed Fischer’s exact test between the difference in total CO versus NCO numbers in mutants from wild-type flies (ns p>0.0083, *p<0.0083, **p<0.0017, ***p<0.00017 after correction for multiple comparisons). Complete dataset can be found in Table S1. **(C)** Mathematical definition of the CE value and table containing centromere number as well as chromosome *2* CE values of WT and *C(1)/0*; *C(4)/0* flies. Two-tailed Fischer’s test was used to calculate significance between observed and expected proximal CO values and was non-significant between *C(1)/0*; *C(4)/0* and wild type flies.

To confirm this reduced centromere number, we performed CENP-C immunofluorescent staining on mitotic chromosome spreads of flies with attached *X* and *4th* chromosomes. CENP-C is a part of the inner kinetochore and is widely used as a centromeric mark in chromosome spreads instead of CENP-A (PALLADINO *et al*. 2020), as the acetic acid fixative required for these spreads tends to degrade histones. Figure 1B shows larval neuroblast spreads and the 12 CENP-C foci we observed in the stock with compound *X* and *4*. This is a 25% reduction in centromere number from wild-type flies, where 16 CENP-C foci were observed (Fig. 1A).

To test whether a reduced centromere number in the *X* and *4th* chromosomes has a *trans*-effect on proximal CO frequencies, we mapped recombination on chromosome *2* through phenotypic scoring of recessive markers. CO distribution between the loci *net* (at the distal end of chromosome *2L*) and *cinnabar* (situated 7.7 Mb into the assembled portion of chromosome *2R*) was measured using flies heterozygous for these markers. This covers a total of 31 Mb on chromosome *2*, with the centromere located in the 11.2 Mb long interval between markers *purple* and *cinnabar,* based on release 6.53 of the *D*. *melanogaster* reference genome.

Fig. 1B shows CO density in the *C(1)/0*; *C(4)/0* mutant plotted across the 31 Mb long region of chromosome *2,* divided by recessive markers into five intervals. We were surprised to observe no change in CO frequencies within the *purple* – *cinnabar* interval which contains the centromere. Reducing centromere number from 16 to 12 did not produce a change in the density of proximal COs, implying that a reduced number of total centromeres does not exert a *trans*-effect on centromere-proximal CO suppression.

Since a distribution graph only compares observed CO frequencies in each interval between mutants and wild-type controls, we sought to also calculate a more biologically relevant measure of CE strength that considers expected versus observed outcomes. This CE value acknowledges that an interval’s expected CO numbers will depend on a genotype’s total CO numbers across the chromosome, thereby accounting for differences in overall CO frequencies between genotypes (Fig. 1C). It allows us to compare observed CO frequencies to those expected were the CE not regulating recombination rates near the centromere. When calculated for *C(1)/0*; *C(4)/0* flies, the CE value on chromosome *2* was 0.90 (Fig. 1C). This is not significantly different from the wild-type CE value of 0.92 on chromosome *2*, further suggesting that dosage changes in centromere number do not have a *trans*-influence on the strength of the centromere effect.

### Increases in total repetitive DNA content do not exert *trans*-effects on the CE

Pericentromeric heterochromatin and repetitive DNA can be thought of as exerting *cis*-effects on CO frequencies in two ways: First, through CO exclusion in highly-repetitive, heterochromatic regions [(HARTMANN *et al*. 2019), (WESTPHAL AND REUTER 2002), (MEHROTRA AND MCKIM 2006; PENG AND KARPEN 2009)], and second, through suppressing COs in adjacent euchromatic, non-repetitive regions [(SLATIS 1955), (JOHN 1985)]. In addition to *cis*-effects, repetitive DNA has also been shown in *D. melanogaster* to influence phenomena such as gene expression and heterochromatin integrity in *trans* [(DIMITRI AND PISANO 1989), (BROWN *et al*. 2020), (DELANOUE *et al*. 2023)]. However, whether repetitive DNA and heterochromatin exert *trans*-effects on centromere-proximal CO frequencies is a question that remains unanswered.

To answer this question, we made use of flies with increased total amounts of repetitive DNA, starting with *XXY,* triplo-*4* flies. The *D. melanogaster* chromosome *4* also consists almost entirely of repetitive sequence and based on chromosome lengths and heterochromatic size estimates from the sequenced genome (HOSKINS *et al*. 2002), the extra *Y* and *4th* chromosomes in an *XXY*, triplo-*4* fly contribute to an ∼37% increase in repetitive DNA as compared to a wild-type *XX* fly (Fig. 2A, C). Similarly, *XXY* flies have an ∼35% increase in repetitive DNA, and triplo-*4* flies an ∼3% increase (Fig. 2B, D). We ensured these mutants did not have increased centromere numbers by using previously validated (Fig. 1A) fly stocks with compound chromosomes of *X* and *4* to build them.

**Figure 2.**
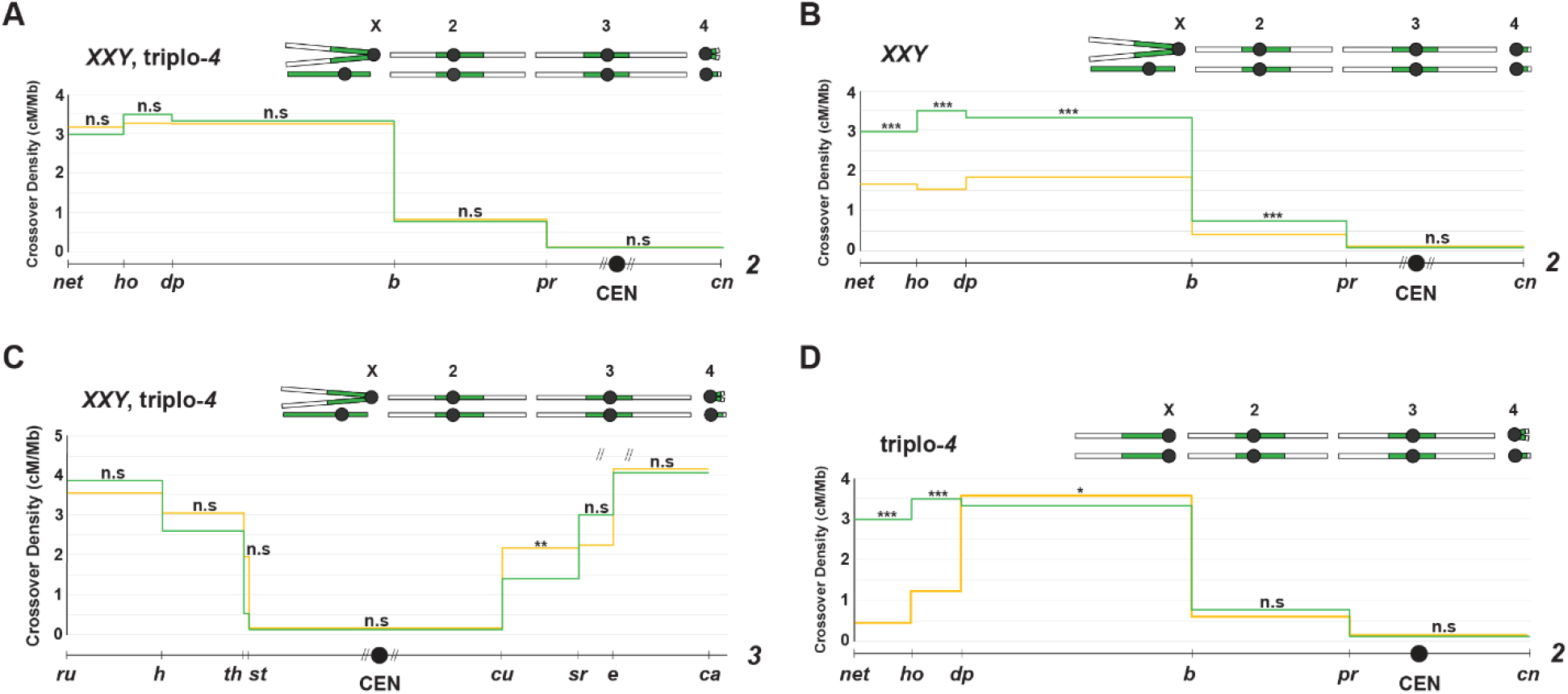
Increases in total repetitive DNA content do not exert *trans*-effects on the CE. **(A)** CO distribution along chromosome *2* in *XXY*, triplo-*4* flies (yellow line, *n*=4434) and wild type flies (green line, *n*= 4104), with a schematic of the *XXY*, triplo-*4* karyotype shown above the graph. CO density in cM/Mb is indicated on the Y-axis, relative physical distances between markers used to score COs are indicated on the X-axis. The black circle represents the centromere, with dashed lines around it representing pericentromeric repetitive sequence that remains unassembled. Statistical significance in each interval was calculated using two-tailed Fischer’s exact test between the difference in total CO versus NCO numbers in mutants from wild-type flies (ns p>0.0083, *p<0.0083, **p<0.0017, ***p<0.00017 after correction for multiple comparisons). Complete dataset can be found in Tables S1. **(B)** CO distribution along chromosome *2* in *XXY* flies (yellow line, *n*=3787) and wild type flies (green line, *n*= 4104), with a schematic of the *XXY* karyotype shown above the graph. Statistical significance in each interval was calculated using two-tailed Fischer’s exact test between the difference in total CO versus NCO numbers in mutants from wild-type flies (ns p>0.0083, *p<0.0083, **p<0.0017, ***p<0.00017 after correction for multiple comparisons). Complete dataset can be found in Tables S1. **(C)** CO distribution along chromosome *3* in *XXY*, triplo-*4* flies (yellow line, *n*=922) and wild type flies (green line, *n*= 1728), with a schematic of the *XXY*, triplo-*4* karyotype shown above the graph. Statistical significance in each interval was calculated using two-tailed Fischer’s exact test between the difference in total CO versus NCO numbers in mutants from wild-type flies (ns p>0.0125, *p<0.0125, **p<0.0025, ***p<0.00025 after correction for multiple comparisons). Complete dataset can be found in Tables S2. **(D)** CO distribution along chromosome *2* in triplo-*4* flies (yellow line, *n*=2924) and wild type flies (green line, *n*= 4104), with a schematic of the triplo-*4* karyotype shown above the graph. Statistical significance in each interval was calculated using two-tailed Fischer’s exact test between the difference in total CO versus NCO numbers in mutants from wild-type flies (ns p>0.0083, *p<0.0083, **p<0.0017, ***p<0.00017 after correction for multiple comparisons). Complete dataset can be found in Tables S1.

To test *trans*-effects of increased repetitive DNA on centromere-proximal CO frequencies, we measured CO distribution across chromosomes *2* and *3* in these mutants, hypothesizing that the decreased heterochromatic integrity in *XXY* and *XYY* flies observed by (BROWN *et al*. 2020) would allow for greater centromere-proximal COs/a weaker CE. Fig. 2B shows CO density plotted across the 31 Mb between markers *net* and *cinnabar* on chromosome *2* in the *XXY*, triplo-*4* mutant. Despite (BROWN *et al*. 2020) observing decreases in canonical heterochromatic marks across the chromosome *2* pericentromere in *XXY* flies, we were surprised to observe no change in centromere-proximal CO frequencies in *XXY*, triplo-*4* flies, compared to wild-type (Fig. 2A). The CE value for this mutant was 0.90, a non-significant difference from the wild-type CE value of 0.92 for chromosome *2* (Table 1).

**Table 1.** CE values in mutants with dosage changes in repetitive DNA content. Table containing estimated changes in repetitive DNA content from wild type (calculated from genome and heterochromatin sizes reported in (HOSKINS *et al*. 2002)) as well as CE values on chromosomes *2* and *3* for various mutants. Two-tailed Fischer’s test was used to calculate significance between observed and expected proximal CO values (** in *XXY* flies and ns for all other mutants, where ns p>0.0083, *p<0.0083, **p<0.0017, ***p<0.00017 on chromosome *2* and ns p>0.0125, *p<0.0125, **p<0.0025, ***p<0.00025 on chromosome *3* after correction for multiple comparisons). Complete datasets can be found in Tables S1 and S2. *Blm* data is from (HATKEVICH *et al*. 2017) with two-tailed Fischer’s test showing extremely (****) significant differences from wild type in observed versus expected proximal CO values (ns p>0.05, *p<0.05, **p<0.01, ***p<0.001, ****p<0.0001).

| Genotype | Estimated change in repetitive DNA content from WT | Chr. 2 CE value | Chr. 3 CE value |
| --- | --- | --- | --- |
| WT | - | 0.92 | 0.90 |
| XXY, triplo-4 | 37% | 0.90 | 0.91 |
| XXY | 35% | 0.84 | 0.85 |
| Triplo-4 | 3% | 0.89 | - |
| *C(1)DX/Y* | <35% | 0.88 | - |
| *Df(2R)M41A10* | 9% | - | 0.87 |
| *Blm* | - | 0.36 | - |

We also measured CO density between the same chromosome *2* markers in flies that were only *XXY*, without an extra copy of chromosome *4.* Here too, we did not observe significant differences in proximal CO frequencies from wild type (Fig. 2C). The CE value for this mutant was 0.84 (Table 1), and although this was moderately significantly different from the wild-type value of 0.92 according to a two-tailed Fischer’s exact text between observed and expected proximal CO values, we do not think this is indicative of a biologically relevant decrease in CE strength for two reasons: 1. On chromosome *2,* the CE value for Blm syndrome helicase mutants, a positive control for CE loss, is 0.36 and is extremely significantly different from wild-type (HATKEVICH *et al*. 2017). The *XXY* CE value is much closer to the wild-type value than that of *Blm* mutants, 2. COs are strongly decreased in this mutant chromosome-wide, with this perceived difference in CE value likely arising from the as yet unknown underlying cause of that.

While we would have liked to measure CO distribution in mutants with a broader range of increases in repetitive DNA content, it is hard to find or build fly stocks that tolerate the structural rearrangements that would make this possible. Due to this limitation, while we have data on the *trans*-effects of larger (∼35 to 37%) increases in repetitive DNA content, we only have one karyotype with an intermediate increase in satellite DNA: *C(1)DX/Y*, another *XXY* mutant with attached *X* chromosomes. Although *C(1)DX/Y* has a *Y* chromosome’s worth of increase in repetitive DNA when compared to *XX* females, the compound *X* chromosomes are missing large regions of pericentromeric heterochromatin, particularly the *rDNA* locus (LINDSLEY AND ZIMM 1992). *C(1)DX* females are always *XXY* as they require *rDNA* on the *Y* chromosome for survival, and at maximum have a ∼30% increase in repetitive DNA content compared to wild-type levels. When we measured CO distribution along chromosome *2* in this mutant, we once again observed no change in CE values compared to wild type (Fig. S1, Table 1).

We also measured centromere-proximal CO frequencies in triplo-*4* flies, which have low (∼3%) increase in repetitive DNA content. These flies were also built using stocks with attached *4th* chromosomes to ensure wild-type centromere numbers. Consistent with our previous data, CO distribution along chromosome *2* in triplo-*4* flies showed no change in centromere-proximal CO frequencies (Fig. 2D) or CE value from wild-type (Table 1), suggesting that no amount of increase in total repetitive DNA content has *trans*-effects on centromere-proximal CO frequencies, or the strength of the centromere effect.

We next investigated whether *XXY* and *XXY*, triplo-*4* mutants have a similar lack of effect on centromere-proximal CO frequencies on other chromosomes as well. To test this, we measured CO distribution along 57.7 Mb of chromosome *3*, between recessive markers *roughoid* and *claret.* The centromere of chromosome *3* lies in the interval between markers *scarlet* and *curled* and contains about 22.8 Mb of assembled sequence. Our chromosome *3* results for *XXY*, triplo-*4* flies followed the same pattern as on chromosome *2,* with no change in proximal CO frequencies, and a CE value of 0.91, not significantly different from the wild-type CE value of 0.90 on chromosome *3* (Fig. 2C). *XXY* flies also followed chromosome *2* patterns, with a CE value of 0.85 (Fig. S1, Table 1), mildly significantly different from the wild-type value of 0.90. However, since this number is much closer to the wild-type range than the CE value of *Blm^-/-^* mutants on chromosome *2* had been, we are once again skeptical of assigning biological relevance to the difference observed. Overall, these results suggest that increasing total repetitive DNA content up to ∼37% has no effect on centromere-proximal CO frequencies in *trans*.

### Decreases in total repetitive DNA content do not exert *trans*-effects on the CE

Lastly, we asked whether decreases in repetitive DNA content can affect CE strength in *trans*. While deleting large chunks of satellite DNA is difficult to do, particularly as large parts of the *D. melanogaster* pericentromere remain unassembled, we were able to use an existing mutant with a large deficiency in chromosome *2, Df(2Rh)M41A10.* This stock has ∼11 Mb of pericentromeric repetitive DNA deleted on chromosome *2* (HILLIKER 1976), which is an ∼9% decrease in total repetitive DNA content in heterozygotes, based on the size of *D. melanogaster* chromosomes and chromatin domains (HOSKINS *et al*. 2002). We procured this mutant from the Bloomington Stock Center, and genetically confirmed the deficiency by crossing to recessive mutants of homozygous lethal genes (*rolled*, *uex*, *Nipped-B*) located within the deleted area.

As *Df(2Rh)M41A10* is on chromosome *2* and does not live as a homozygote, we measured CO distribution along chromosome *3* in heterozygotes to test whether a decrease in repetitive DNA content influences proximal CO frequencies in *trans*. Consistent with our previous data, we observed no change in proximal CO numbers (Fig. 3) or CE value (Table 1), leading us to strongly conclude that, as with centromere number, dosage changes in repetitive DNA content are unable to exert a *trans*-effect on the centromere effect.

**Figure 3.**
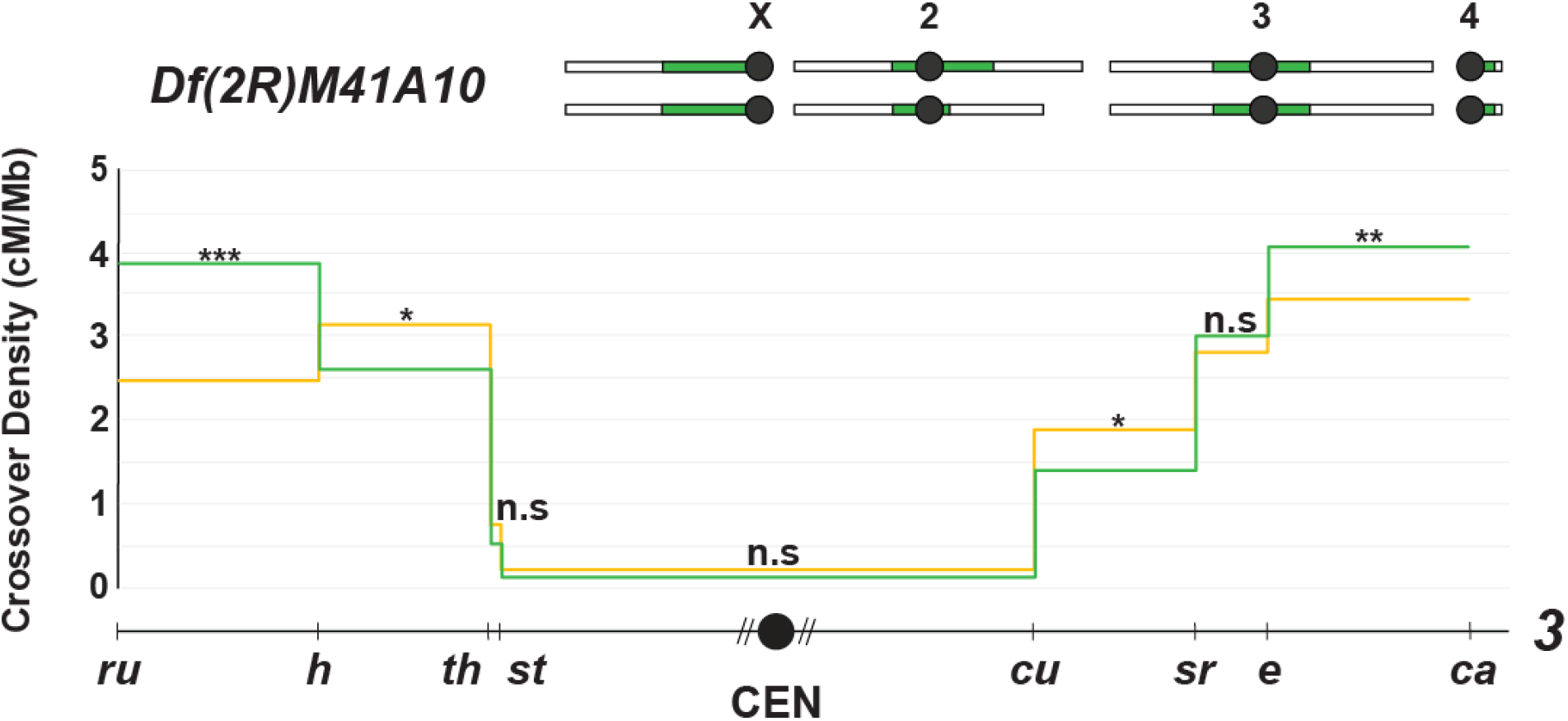
Decreases in total repetitive DNA content do not exert *trans*-effects on the CE. CO distribution along chromosome *3* in *Df(2R)M41A10* flies (yellow line, *n*=1773) and wild type flies (green line, *n*= 1728), with a schematic of the *Df(2R)M41A10* karyotype shown above the graph. CO density in cM/Mb is indicated on the Y-axis, relative physical distances between markers used to score COs are indicated on the X-axis. The black circle represents the centromere, with dashed lines around it representing pericentromeric repetitive sequence that remains unassembled. Statistical significance in each interval was calculated using two-tailed Fischer’s exact test between the difference in total CO versus NCO numbers in mutants from wild-type flies (ns p>0.0125, *p<0.0125, **p<0.0025, ***p<0.00025 after correction for multiple comparisons). Complete dataset can be found in Tables S2.

### No *trans*-effects of satellite DNA-binding proteins on the CE

On observing that satellite DNA dosage does not have *trans*-effects on CE strength, we wondered whether this was also true of satellite DNA function. Although historically thought of as genomic junk, we now know that satellite DNA is important in many ways: protecting centromeric regions from evolutionary forces (YUNIS AND YASMINEH 1971), forming and structurally defining centromeric chromatin (MURPHY AND KARPEN 1998), acting as a fertility barrier between species [(FERREE AND BARBASH 2009), (JAGANNATHAN AND YAMASHITA 2021)], and facilitating chromosome pairing and segregation of achiasmate chromosomes (DERNBURG *et al*. 1996). We sought to determine whether satellite DNA function is important for the manifestation of the CE by measuring centromere-proximal CO frequencies in mutants of *D*. *melanogaster* satellite DNA-binding proteins D1 and Proliferation disrupter (Prod).

D1 is an AT-hook protein that binds to {AATAT}_n_ satellite, while Prod binds to Prodsat {AATAACATAG}_n_. The satellite DNA-binding functions of D1 and Prod were shown recently to be important for chromocenter formation in *D*. *melanogaster* spermatocytes and imaginal discs, respectively [(JAGANNATHAN *et al*. 2018), (JAGANNATHAN *et al*. 2019)]. Since our previous results indicate that dosage of pericentromeric repetitive DNA has no *trans*-effect on the CE, we asked whether proteins known to be important for satellite DNA function are involved in CE manifestation, by looking at CO formation and CE strength in *D1* and *Prod* mutants.

Fig. 4A shows CO distribution along chromosome *2* in *D1^LL03310^/Df(3R)BSC^666^,* a genotype shown to result in de-clustering of {AATAT}_n_ satellite DNA (JAGANNATHAN *et al*. 2018). We observed no significant change in CO frequencies in the centromeric interval, with CE value being 0.90, a non-significant change from the chromosome *2* CE value of 0.92 in wild-type flies. We also measured CO distribution along chromosome *3* in *Prod^K^/+* mutants and observed no change in centromere-proximal CO frequencies or CE value when compared to wild-type (Fig. 4B, C). However, it must be noted that *Prod* is an essential gene and since homozygous mutants are inviable, we were limited to measuring recombination in females heterozygous for a null mutation. Collectively, these results suggest that satellite DNA-binding proteins do not have a *trans*-effect in manifesting the CE, a pattern consistent with what we see for satellite DNA dosage.

**Figure 4.**
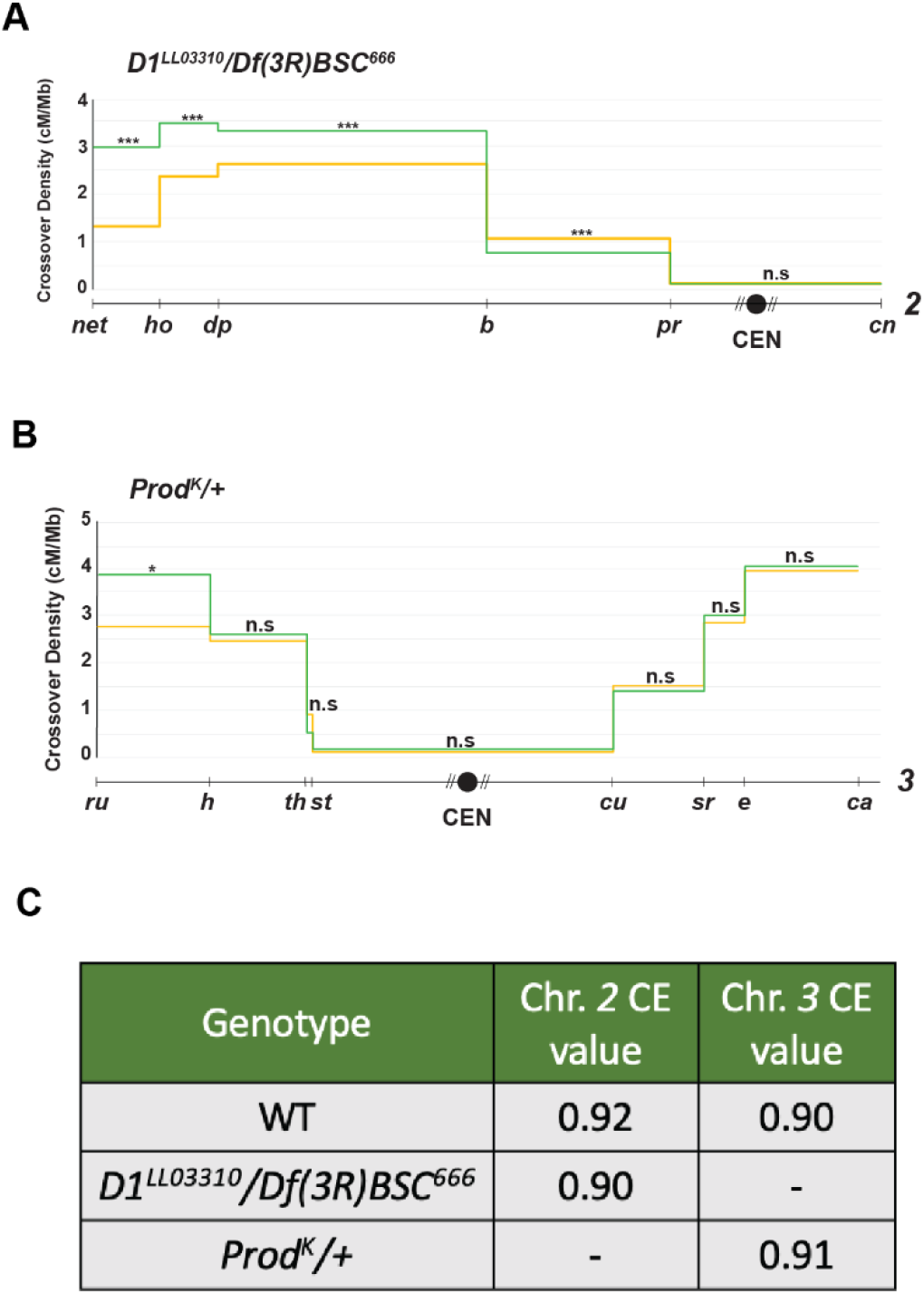
No *trans*-effects of satellite DNA-binding proteins on the CE. **(A)** CO distribution along chromosome *2* in *D1^LL03310^*/*Df(3R)BSC^666^* flies (yellow line, *n*=6399) and wild type flies (green line, *n*= 4104). CO density in cM/Mb is indicated on the Y-axis, relative physical distances between markers used to score COs are indicated on the X-axis. The black circle represents the centromere, with dashed lines around it representing pericentromeric repetitive sequence that remains unassembled. Statistical significance in each interval was calculated using two-tailed Fischer’s exact test between the difference in total CO versus NCO numbers in mutants from wild-type flies (ns p>0.0083, *p<0.0083, **p<0.0017, ***p<0.00017 after correction for multiple comparisons). Complete dataset can be found in Tables S1. **(B)** CO distribution along chromosome *3* in *Prod^K^/+* flies (yellow line, *n*=5320) and wild type flies (green line, *n*= 1728). Statistical significance in each interval was calculated using two-tailed Fischer’s exact test between the difference in total CO versus NCO numbers in mutants from wild-type flies (ns p>0.0125, *p<0.0125, **p<0.0025, ***p<0.00025 after correction for multiple comparisons). Complete dataset can be found in Tables S2. **(C)** Table containing CE values on chromosomes *2* and *3* of *D1^LL03310^*/*Df(3R)BSC^666^* and *Prod^K^/+* flies, respectively. Two-tailed Fischer’s test was used to calculate significance between observed and expected proximal CO values and was non-significantly different from wild type for both mutants.

## Discussion

Studies on the *D. melanogaster* CE have historically focused on the local, *cis*-acting contributions of the centromere and pericentromeric heterochromatin. In this study, we test whether dosage changes in certain structural components of chromosomes - centromeres and repetitive DNA - exert *trans*-effects on centromere-proximal CO suppression in *D. melanogaster*. Despite previous studies suggesting that both factors are likely to have genome-wide effects on this patterning phenomenon, our study finds that the CE is surprisingly robust to dosage changes in both centromere number and quantity of repetitive DNA. Below, we discuss the mechanistic interpretations of these findings.

### Centromere number and the CE

On reducing total centromere number using fly stocks with compound chromosomes of *X* and *4*, we expected to see a reduction in centromere-proximal CO frequencies, based on Redfield’s observation of increased proximal-crossovers in triploids (that have 24 centromeres compared to 16 in diploids) [(REDFIELD 1930), (REDFIELD 1932)]. This would suggest that centromeres can act as molecular sinks to a putative “CE factor” necessary to maintain wild-type levels of centromere-proximal CO suppression, with a dosage reduction in centromere number leading to increased availability of this “CE factor”, potentially leading to a strengthened CE. Alternatively, increased centromere-proximal CO in flies with a reduced number of centromeres would suggest that a baseline centromere number is required to maintain the CE at wild-type levels. With centromere clustering observed during meiotic prophase I when recombination occurs (HATKEVICH *et al*. 2021), this would potentially have indicated a role for the chromocenter in establishing the CE.

However, in our study we observed no change in CE strength when total centromere number is reduced, suggesting that centromeres are neither acting as molecular sinks nor spatially regulating centromere-proximal CO frequencies within the chromocenter. This deviation from what we expected based on Redfield’s data suggests that the increase observed in her study is likely not due to changes in total centromere number, and perhaps a result of ploidy changes in triploids. It is also possible that changing centromere numbers could only be exerting *cis*-effects on centromere-proximal COs, a hypothesis that is hard to test as mapping recombination within compound chromosomes is near impossible. Another possibility is that reducing total centromere number by 4 is not sufficient to cause an effect. To reduce centromere numbers further, we would need mutants with an even greater number of attached chromosomes. Unfortunately, flies heterozygous for markers in this background are highly difficult to generate, if not inviable.

### Repetitive DNA content and the CE

Westphal and Reuter demonstrated in 2002 that mutants of *Su(var)* genes coding for important heterochromatin factors such as HP1 and H3K9 methyltransferase displayed an increase in centromere-proximal CO frequencies (WESTPHAL AND REUTER 2002). Combined with extensive evidence that extra copies of chromosome *Y* affect pericentromeric heterochromatinization genome-wide in *D. melanogaster* [(DIMITRI AND PISANO 1989), (BROWN *et al*. 2020), (DELANOUE *et al*. 2023)], we hypothesized that centromere-proximal CO frequencies would be increased in *XXY* flies and other mutants with dosage changes in repetitive DNA content. This would be consistent with extra repetitive DNA soaking up factors necessary to establish and maintain pericentromeric heterochromatin/the CE, leading to decreased availability in the rest of the genome, potentially allowing for increased centromere-proximal COs. Alternatively, increased repetitive DNA may have led to a stronger CE, which would suggest a more direct role for repetitive DNA in maintaining CO suppression around the centromere.

In our study we observed no change in centromere-proximal CO frequencies with dosage changes in repetitive DNA, suggesting that the loss of heterochromatinization observed in *XXY* flies by (BROWN *et al*. 2020) is not sufficient to allow for increased centromere-proximal COs. The increase in proximal COs observed by (WESTPHAL AND REUTER 2002) in certain *Su(var)* mutants with defective heterochromatinization suggests that there is a threshold of heterochromatin loss necessary for an increase in CO frequencies; if true, our study indicates that *XXY* a 37% increase in repetitive DNA content do not reach this threshold. Interestingly, the phenomenon of position effect variegation in *XXY* flies suggests that the threshold of heterochromatic disruption necessary for changes in gene expression is lower than that necessary for recombination, and that the open chromatin landscape in these mutant flies can be permissive to transcriptional machinery without being permissive to DSB/recombination machinery. Additionally, the lack of change observed in our study also rules out spatial regulation of the CE by repetitive DNA - potentially within structures such as the chromocenter - as dosage changes in repetitive DNA content were not directly proportional to CE strength.

### Repetitive DNA function and the CE

As our study demonstrates that the *D. melanogaster* CE is unaffected by dosage changes in repetitive DNA, we wondered whether disrupting repetitive DNA function using satellite DNA binding-protein mutants would have genome-wide effects on the CE. To test this, we measured centromere-proximal CO frequencies in mutant that lack the AT-hook protein D1, which binds to {AATAT}_n_ satellite and has been shown by (JAGANNATHAN *et al*. 2018) to disrupt chromocenters. Surprisingly, we once again observed no change in CE strength in this, or a *Prod* mutant, suggesting that satellite DNA-binding proteins do not play a role in establishing the CE in *D. melanogaster*.

While this result could be interpreted as {AATAT}_n_ satellite clustering/chromocenters not being necessary to maintain CO suppression around the centromere, we hesitate to strongly state this conclusion as it is possible that the declustering observed in fly spermatocytes by (JAGANNATHAN *et al*. 2018) may not happen to the same extent in oocytes, although centromere clustering *is* widely observed in the female germline during meiotic prophase I and exonic expression of D1 occurs at the same levels in both males and females (GRAMATES *et al*. 2022). It is also noteworthy that the {AATAT}_n_ satellite is primarily found on the *X* chromosome, making it possible that CE disruption in D1 mutants only takes place in *cis*. This can be tested by measuring CE strength on chromosome *X* in these mutants, although the CE is already very weak on this chromosome, likely due to the expanse of satellite DNA on it. Despite these caveats, our results do indeed demonstrate that the {AATAT}_n_ satellite binding protein D1 does not have global roles in establishing the *D. melanogaster* CE.

### CE mechanism

Our study rules out the mechanistic role of structural chromosomal components such as centromeres and repetitive DNA in suppressing centromere-proximal COs in *trans*, suggesting that the *D. melanogaster* CE is more likely to be genetically than spatially controlled. Although previous studies from our lab have shown that CO interference and assurance are genetically separable from the CE (BRADY *et al*. 2018), the emergence of interference models such as coarsening makes us question whether the CE could also be explained by variations of these models. Post-translational modifications of pro-crossover meiotic proteins are being heavily investigated for their role in establishing CO interference [(HAVERSAT *et al*. 2022), (ZHANG *et al*. 2021)], and a speculative idea we would like to put forth is that such modifications could also be manifesting the CE through modifying enzymes like kinases being anchored at the centromere and biasing neighboring chromosomal regions - less-repetitive beta heterochromatin and proximal euchromatin - towards NCO repair. While the factors involved may be different from those in CO interference, such a model sees both patterning phenomena manifesting through similar modes of genetic control, an idea that is further supported by our study ruling out structural and spatial contributions to centromere-proximal CO suppression.

## Conclusions

In conclusion, our study shows that the CE is robust to dosage changes in certain structural components of chromosomes, such as centromeres and repetitive DNA. This suggests that the CE is likely not controlled spatially during meiotic prophase and is perhaps mediated through genetic factors, opening up avenues of future research to uncover the exact mechanistic details of centromere-proximal CO suppression in *D. melanogaster*.

## Materials and Methods

### *Drosophila* stocks

Flies were maintained on standard medium at 25C. The following fly strains were obtained from the Bloomington Stock Center: 1612 (*C(1)RM, y^1^ v^1^/C(1;Y)^1^, v^1^ f^1^ Bar^1^/0; C(4)RM, ci^1^ ey^R^/0*), 9460 (*C(1)RM/C(1;Y)6, y^1^ w^1^ f^1^/0)*, 1785 *(C(4)RM, ci^1^ ey^R^/0),* 741 (*Df(2R)M41A10/SM1*). The following fly strains were generously gifted to us by Dr. Yukiko Yamashita: *Prod^K^/CyO, Act::GFP*, *w*; *Df(3R)BSC^666^/TM6C, Sb*, *D1^LL03310^*/*TM6B, Tb.* Wild type controls were *Oregon-R* which was gifted to us by Dr. Scott Hawley.

### Genetic assays

Chromosome *2* crossovers were mapped by crossing virgin *net dpp^ho^ dp b pr cn* /+ females of desired mutant backgrounds to males that were homozygous *net dpp^ho^ dp b pr cn*. Chromosome *3* crossovers were mapped by crossing virgin *ru h th st cu sr e ca* /+ females of desired mutant backgrounds to males that were homozygous *ru h th st cu sr e ca.* Vials were set up with 1-5 day old females, then flipped a week later. For both chromosomes *2* and *3*, progeny were scored for all markers.

### Fly crosses

Compound females (*C(1)RM, y^1^ v^1^/0*; *C(4)RM, ci^1^ ey^R^/0*, *C(1)RM/0*, *(C(4)RM, ci^1^ ey^R^/0*) were crossed to males homozygous for recessive markers on chromosomes *2* or *3* to obtain *XXY*, triplo-*4* (*C(1)RM, y^1^ v^1^/Y*; *C(4)RM, ci^1^ ey^R^ /+*), *XXY* (*C(1)RM/Y*), and triplo-*4* (*C(4)RM, ci^1^ ey^R^/+*) females heterozygous for recessive markers.

### Recombination calculation

Genetic distance is calculated as 100 * (r/n) where n is total scored progeny and r is total recombinant progeny within an interval (including single, double, and triple crossovers). It is expressed in centiMorgans (cM). Variance was used to calculate 95% confidence intervals, as in (STEVENS 1936). Physical distances between recessive markers used for phenotypic crossover mapping were calculated using positions of genetic markers on release 6.53 of the *Drosophila melanogaster* reference genome. Distances were calculated from the beginning of a genetic marker to the end of the previous marker. CE values were calculated as 1 - (observed COs/expected COs), where expected COs are calculated as follows: total COs * (length of proximal interval/total length).

### Larval neuroblast chromosome spreads

Brains were dissected from wandering third instar larvae in cold PBS, incubated in 0.5% sodium citrate for 10 minutes, then fixed in 2% formaldehyde and 45% acetic acid solution for 7 minutes on siliconized cover-slips. Brains were squashed onto glass slides (VWR Micro Slides, Superfrost Plus) and frozen in liquid nitrogen, then washed thrice in PBS-T for 10 minutes each. Slides were blocked for 1 hour in 1% BSA, PBS-T solution, incubated overnight with anti-CENP-C antibody (1:5000 from Dr. Kim McKim) at 4C. They were then washed four times in PBS-T for 10 minutes each and incubated for 2 hours in secondary antibody (1:500, anti-rabbit) at room temperature. Slides were washed again in PBS-T, four times for 10 minutes each, then mounted with a 1:1000 solution of 1 mg/ml DAPI in Fluoromount-G.

### Imaging

Chromosome spreads of larval brains were imaged using a Zeiss LSM880 confocal laser scanning microscope with Airyscan, using the 63x oil-immersion objective lens. FIJI (ImageJ) was used to process images.

## Data Availability Statement

*Drosophila* stocks are available upon request. The authors confirm that all data necessary for confirming the conclusions of the article are present within the article, figures, table, and supplemental information.

## Acknowledgments

We thank Yukiko Yamashita for the fly stocks that were generously sent to us, Yves Barral for the suggestion that a centromere-anchored kinase might contribute to the CE, and members of the Sekelsky lab for helpful comments on the manuscript.

## Funding

This work was supported by a grant from the National Institute of General Medical Sciences to JS under award 1R35GM118127. NP was supported in part by a grant from the National Institute on Aging under award 1F31AG079626-01 and the National Institute of General Medical Sciences under award T32GM135128. LF was supported in part by a grant from the National Science Foundation Division of Biological Infrastructure under award 2048087.

## Competing Interests

The authors declare that they have no conflicts of interest.

---

**<u>Figure S1.</u> (A)** Mitotic chromosome spreads (grey) with CENP-C foci (magenta) from *C(1)/0* larvae (top panel) and wild type larvae (bottom panel). Green boxes highlight chromosome *X* (larger, V-shaped structure) in both genotypes. **(B)** CO distribution along chromosome *2* in *C(1)DX/Y* flies (yellow line, *n*=2702) and wild type flies (green line, *n*= 4104). CO density in cM/Mb is indicated on the Y-axis, relative physical distances between markers used to score COs are indicated on the X-axis. The black circle represents the centromere, with dashed lines around it representing pericentromeric repetitive sequence that remains unassembled.

Statistical significance in each interval was calculated using two-tailed Fischer’s exact test between the difference in total CO versus NCO numbers in mutants from wild-type flies (ns p>0.0083, *p<0.0083, **p<0.0017, ***p<0.00017 after correction for multiple comparisons). Complete dataset can be found in Table S1. **(C)** CO distribution along chromosome *3* in *C(1)RM/Y* flies (yellow line, *n*=1854) and wild type flies (green line, *n*= 1728). CO density in cM/Mb is indicated on the Y-axis, relative physical distances between markers used to score COs are indicated on the X-axis. The black circle represents the centromere, with dashed lines around it representing pericentromeric repetitive sequence that remains unassembled.

Statistical significance in each interval was calculated using two-tailed Fischer’s exact test between the difference in total CO versus NCO numbers in mutants from wild-type flies (ns p>0.0125, *p<0.0125, **p<0.0025, ***p<0.00025 after correction for multiple comparisons). Complete dataset can be found in Tables S2.

**<u>Table S1.</u>** Table showing the complete dataset used to make the chromosome *2* graphs in Fig. 1, 2, 4, and S1. The number of parental, single crossover (SCO), double crossover (DCO), and triple crossover (TCO) flies in each mutant are shown.

**<u>Table S2.</u>** Table showing the complete dataset used to make the chromosome *3* graphs in Fig. 2, 3, 4, and S1. The number of parental, single crossover (SCO), double crossover (DCO), and triple crossover (TCO) flies in each mutant are shown.

